# Surprisingly weak coordination between leaf structure and function among closely-related tomato species

**DOI:** 10.1101/031328

**Authors:** Christopher D. Muir, Miquel À. Conesa, Emilio J. Roldán, Arántzazu Molins, Jeroni Galmés

**Affiliations:** Department of Biology, Indiana University, Bloomington, IN 47405, USA; Biodiversity Research Centre and Botany Department, University of British Columbia, Vancouver, British Columbia V6T 1Z4, Canada; Research Group on Plant Biology under Mediterranean Conditions, Departament de Biologia, Universitat de les Illes Balears, Ctra. Valldemossa km 7.5, E-07122, Palma de Mallorca, Spain.

## Abstract

Natural selection may often favor coordination between different traits, or phenotypic integration, in order to most efficiently acquire and deploy scarce resources. As leaves are the primary photosynthetic organ in plants, many have proposed that leaf physiology, biochemistry, and anatomical structure are coordinated along a functional trait spectrum from fast, resource-acquisitive syndromes to slow, resource-conservative syndromes. However, the coordination hypothesis has rarely been tested at a phylogenetic scale most relevant for understanding rapid adaptation in the recent past or predicting evolutionary trajectories in response to climate change. To that end, we used a common garden to examine genetically-based coordination between leaf traits across 19 wild and cultivated tomato taxa. We found surprisingly weak integration between photosynthetic rate, leaf structure, biochemical capacity, and CO_2_ diffusion, even though all were arrayed in the predicted direction along a ‘fast-slow’ spectrum. This suggests considerable scope for unique trait combinations to evolve in response to new environments or in crop breeding. In particular, we find that partially independent variation in stomatal and mesophyll conductance may allow a plant to improve water-use efficiency without necessarily sacrificing maximum photosynthetic rates. Our study does not imply that functional trait spectra or tradeoffs are unimportant, but that the many important axes of variation within a taxonomic group may be unique and not generalizable to other taxa.

## Introduction

The ecology of organisms critically depends on their ability to obtain energy for growth and reproduction. In C_3_ plants, both passive diffusion of CO_2_ from the atmosphere to chloroplasts, via stomata and the leaf mesophyll, and biochemical capacity limit photosynthetic rates (Farquhar & Sharkey 1982; Parkhurst 1994; Evans *et al*. 2009; Flexas *et al*. 2015). Optimization theory predicts that these functions should be tightly coordinated: leaf anatomies that allow rapid diffusion of CO_2_ should also have the biochemical capacity to take advantage of it and *vice versa* (Givnish 1986; Wright *et al*. 2004; Reich 2014). Natural selection on trait coordination, also known as phenotypic integration (Pigliucci 2003), should thus generate major functional trait spectra associated with different ecological strategies (Grime 1977; Westoby *et al*. 2002; Chave *et al*. 2009; Reich 2014). Indeed, intrinsic photosynthetic capacity varies widely between species, indicating that leaf-level CO_2_ diffusion and biochemistry are major levers through which natural selection and crop breeders can alter plant performance and fitness in different environments. For example, selection might favor leaf traits that inhibit rapid CO_2_ diffusion (e.g. lower stomatal density or thicker leaves) if doing so has an adaptive benefit, bringing about a tradeoff (e.g. sclerophyll leaves of species from water-limited environments [Medrano *et al*. 2009]). It is not clear whether most functional trait variation between species lies along a single major axis (e.g. fast-slow continuum [Reich 2014]) or orthogonally along multiple independent trait axes.

In particular, the presence of functional trait spectra has rarely been examined among closely-related species in a phylogenetic context (Edwards *et al*. 2014; Mason & Donovan 2015). However, for certain ecological questions, data on genetically-based variation among closely-related species, rather than broad comparisons across disparate families and plant functional types, are most appropriate (Donovan *et al*. 2014). Specifically, ecological diversification is not always predictable because different groups of organisms find unique adaptive solutions, leading to multiple trait spectra. For example, the leaf economics spectrum (Wright *et al*. 2004) is not a one-size-fits-all trait axis, but rather a set of axes that vary between communities (Funk & Cornwell 2013) and taxa (Edwards et al. 2014; Mason & Donovan 2015). Second, the evolutionary routes (not) taken in the recent past among closely-related species may help predict how species will respond to natural and anthropogenic climate change (Kellerman *et al*. 2012; Donovan *et al*. 2014). Finally, crop breeders can take advantage of existing trait variation in crop-wild relatives to develop new varieties for sustainable agriculture (Moyle & Muir 2010; Dennison 2012; Giuliani *et al*. 2013). Hence, if coordination among leaf traits is limited among closely-related species, then broad-scale or global trait spectra may be of little use in addressing fundamental challenges like the ecology of local adaptation (Kawecki & Ebert 2004), near-term responses to climate change, and crop breeding. Conversely, weak coordination may indicate significant opportunities for natural and artificial selection to fine-tune leaf traits in response to different ecological circumstances.

We used a common garden experiment to measure coordination between leaf traits, specifically diffusive and biochemical limitations on photosynthesis and water-use efficiency as well as leaf structure. We considered these traits in relation to climate of origin among tomato species (*Solanum* sect. *Lycopersicon*) and their relatives (*Solanum* sect. *Juglandifolia* and *Lycopersicoides*). These species are closely related herbaceous perennials: fossil-calibrated molecular clocks put section *Lycopersicon* at approximately 2.48 my old, and the split between *Lycopersicon* and *Lycopersicoides* at approximately 4.7 my ago (Särkinen *et al*. 2013). Despite sharing a recent common ancestor, tomato species are ecologically and phenotypically diverse (Moyle 2008; Peralta *et al*. 2008; Nakazato *et al*. 2008, 2012; Chitwood *et al*. 2012; Haak *et al*. 2014; Muir & Thomas-Huebner 2015). Adaptation to different climates, especially along temperature and precipitation gradients, may have played an important role in diversification (Nakazato *et al*. 2010). Most wild tomato species are interfertile (e.g. Baek *et al*. 2015) with the domesticated tomato (*S. lycopersicum* var. *esculentum*), providing a valuable source of germplasm for crop improvement. In particular, altering leaf CO_2_ diffusion properties may enhance photosynthetic rate and/or water-use efficiency (Tholen *et al*. 2012; Flexas *et al*. 2013; Gago *et al*. 2014; Buckley & Warren 2014; Flexas *et al*. 2015). Diffusion limits photosynthesis in some wild tomato species (Muir *et al*. 2014) and cultivars (Galmés *et al*. 2011, 2013), but we know little about the ecological and evolutionary significance of this variation.

More generally, we know relatively more about the mechanistic basis of variation in leaf photosynthesis than its ecological or adaptive significance. Stomatal and mesophyll CO_2_ diffusion are strongly affected by leaf anatomy (Brown & Escombe 1900; Franks & Farquhar 2001; Nobel 2009; Tholen and Zhu 2011; Tomás *et al*. 2013); biochemical limitations depend on the amount of Rubisco, its activation and catalytic rates (Farquhar *et al*. 1980; von Caemmerer 2000; Galmés *et al*. 2014). Maximum photosynthetic rates are generally higher among certain plant functional types (herbs, C_3_ graminoids, and crop plants) than others (trees, sclerophylls, and succulents) (Körner *et al*. 1979; Wullschleger 1993; Flexas *et al*. 2008). These broad-scale correlations suggest that slow growing plants tradeoff high photosynthetic rate in favor of leaf traits that may have other fitness benefits, such as durability, defense against herbivores, or increased resource-use efficiency (Turner 1994; Aerts 1995; Onoda *et al*. 2011).

We found surprisingly weak coordination among CO_2_ diffusion, biochemical capacity, photosynthesis, water-use efficiency, and leaf structure among 19 wild and cultivated tomato taxa. The principal axis of variation indicates an expected tradeoff between robust leaf structure and photosynthesis. However, there was substantial variation orthogonal to this axis that allowed some species simultaneously achieve relatively high photosynthetic rates and intrinsic water-use efficiency. Also, contrary to expectations from the literature, there was no indication that more conservative leaf traits were associated with drier or hotter environments. If anything, there was a signature of less robust leaf structure among species from the driest habitats. Thus, our data suggest that functional trait spectra inferred from global comparisons across plant families and functional types may be of limited utility for predicting short-term evolutionary responses.

## Methods

### Plant materials and cultivation

We obtained seeds of 19 taxa, 16 wild species (Table S1) from the Tomato Genetics Resource Center at UC Davis (TGRC; http://tgrc.ucdavis.edu) as well as 3 cultivated accessions of *S. lycopersicum* var. *esculentum*, cv. ‘Roma VF’ (Batlle S.A.) and two Mediterranean ‘Tomàtiga de Ramellet’ accessions from the University of the Balearic Islands seedbank collection (UIB1–30 and UIB1–48). To even out plant size during measurements, and following TGRC indications, slower growing species were germinated two weeks ahead of faster growing species. Seeds were soaked in 2.5% sodium hypochlorite (household bleach) for 30 or 60 minutes (following TGRC instructions for each species), rinsed thoroughly, and placed on moist paper to germinate. After one week, seedlings were transplanted to cell-pack flats. Two weeks later, five plants of each taxa were transplanted to 19 L pots where the experiment was performed. Pots contained a mixture of standard horticultural substrate mixed with pearlite in a 4:1 proportion v/v. Plants were grown outdoors in an open experimental field under typical Mediterranean conditions at the University of the Balearic Islands (39° 38’ 14.9’’ N, 2° 38’ 51.5’’ E) during spring-summer season. Plants were irrigated to field capacity daily to prevent drought stress and fertilized weekly with an NPK solution.

### Diffusional and biochemical constraints on photosynthesis and water-use efficiency

We measured stomatal (*g*_s_), mesophyll (*g*_m_) conductance to CO_2_, net CO_2_ assimilation rate (*A*_N_) at ambient CO_2_ concentrations, and intrinsic water-use efficiency (*iWUE* = *A*_N_ /*g*_sw_) using an open-path infrared gas exchange analyzer (LI-6400 or LI-6400XT, LI-COR Inc., Lincoln, NE, USA) with a 2-cm^2^ leaf chamber fluorometer. Note that *g*_s_ and *g*_sw_ refer to stomatal conductance to CO_2_ and H_2_O, respectively. Each leaf acclimatized in the chamber until steady state (usually 15–30 min) under standardized conditions: ambient CO_2_ (*C*_a_ = 400 ppm); constant leaf temperature (*T*_leaf_ = 25° C), saturating irradiance (photosynthetically active radiation, PAR = 1500 μmol quanta m^−2^ s^−1^), and moderate humidity (relative humidity = 40–60%). We took point measurements of all traits under these steady-state conditions. Additionally, we calculated the maximum rate of carboxylation (*V*_cmax_) using *A-C*_c_ curves and leaf dark respiration (*R*_dark_) at predawn. Estimating *g*_m_ can be particularly sensitive to assumed parameter values. To accurately measure *g*_m_ we estimated species-specific parameters of leaf respiration, light absorptance/photosystem partitioning, and Rubisco kinetic parameters. Further detail on measuring *g*_m_ using combined gas exchange and chlorophyll fluorescence is provided in Methods S1. To investigate Rubisco kinetics, we also sequenced the *rbc*L gene encoding the Rubisco large subunit (LSu) from each species to identify amino acid sequence changes that could alter Rubisco kinetics (see Methods S1 for further detail). Next, we characterized Rubisco kinetic parameters directly from two species (*S. lycopersicum* var. *esculentum*, cv. ‘Roma VF’ and *S. lycopersicoides*) representing the two Rubisco LSu types identified from sequence analysis (Methods S1). Kinetic properties of the two Rubisco LSu types were compared using ANOVA.

### Leaf anatomical measurements

We used the youngest, unshaded, fully-expanded leaf from each individual. Leaves were immediately weighed and scanned to obtain fresh mass (FM) and leaf area (LA), respectively. Afterwards, leaves were dried for at least 48 h in a drying oven at 60 °C to obtain dry mass (DM). We report *LMA* (as DM/LA) on whole leaves, which in tomato are pinnately compound, but we excluded structural, non-laminar portions (petiole, rachis, and petioules) because we were particularly interested tradeoffs between leaf structure and diffusive conductance within the lamina. Because leaf thickness measurements using a micrometer are unreliable in tomato leaves (C.D. Muir, pers. obs), we estimated leaf thickness (*LT*) using the method of Vile *et al*. (2005):

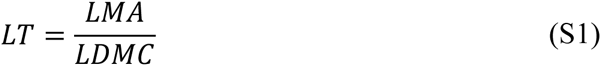

*LDMC* is the leaf dry matter content, the ratio of leaf DM to saturated FM. Leaf thickness calculated using Eq. 1 is closely correlated with leaf thickness measured from sections in tomatoes (Muir *et al*. 2014). We obtained leaf morphological data from 80 of 82 individuals.

### Statistical analyses

We used phylogenetic linear mixed effects models (‘phyloLME’) to estimate key relationships among traits while accounting for phylogenetic nonindependence. Specifically, we fit statistical models using a Bayesian Markov chain Monte Carlo (MCMC) algorithm implemented in the R package MCMCglmm version 2.21 (Hadfield 2010). For all models, we ran the MCMC chain under diffuse priors for 10^6^ steps after a burn-in of 10^5^ steps, sampling the posterior distribution 10^4^ times every 10^2^ steps. We tested whether stomatal and mesophyll conductance were significantly correlated with *A*_N_ and *iWUE*. These variables were log-transformed for linearity and homoscedasticity. We estimated the effect of these diffusion traits on *A*_N_ and *iWUE* from the mode of the posterior distribution and inferred statistical significance if the 95% highest posterior density (HPD) interval did not overlap zero. We simultaneously tested whether phylogeny explained photosynthetic trait variation by including Species as a phylogenetically-structured random effect and compared that to a model without Species using the deviance information criterion (DIC), where a decrease of 2 or more is interpreted as a significant increase in model fit. We used a maximum likelihood phylogenetic tree inferred from 18 genes (Figure 1; Haak *et al*. 2014). Maximum likelihood analyses were conducted using RAxML version 8.1.24 (Stamatakis 2014). The topology of the best tree agreed with previous Bayesian estimates (Rodriguez *et al*. 2009).

**Figure 1:**
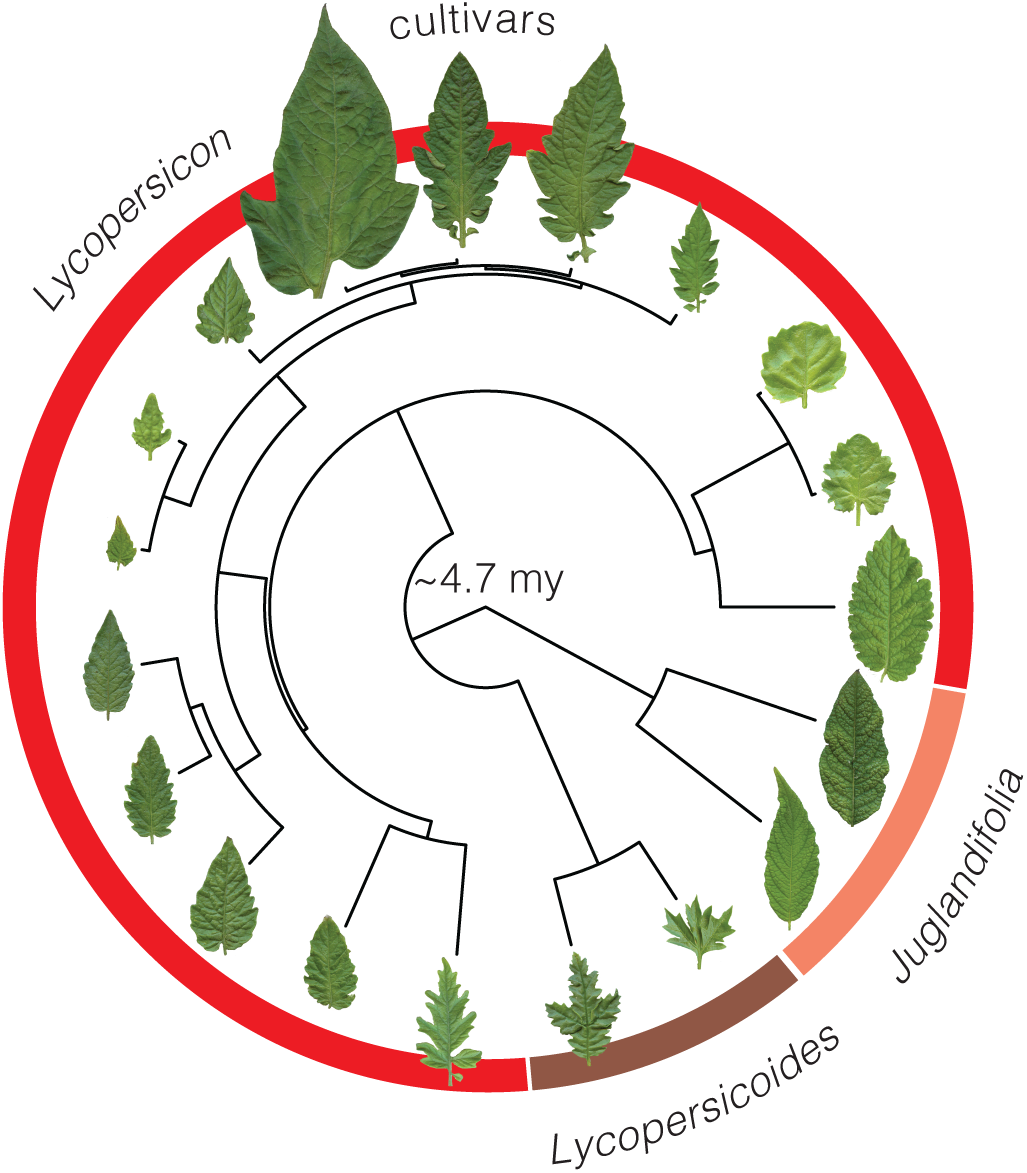
Wild tomatoes and cultivars are closely-related, yet phenotypically-diverse taxa. Among species investigated here, the oldest split is approximately 4.7 my based on a fossil-calibrated phylogeny (Särkinen *et al*. 2013). Functional traits like terminal leaflet size, depicted for each species at the tips of the phylogeny, as well as leaf physiological and structural traits used in this study vary widely among species.

We used principal component analysis (PCA) to identify major axes of variation among nine leaf traits: *g*_m_, *g*_s_, *A*_N_, *V*_cmax_, *R*_dark_, *iWUE, LMA, LT, LDMC*. All traits except *V*_cmax_ and *R*_dark_ were log-transformed to make the distribution approximately multivariate normal. Parallel analysis of the trait correlation matrix using the ‘parallel’ function from the R package nFactors version 2.3.3 (Raiche 2010) indicated that the first four principal components explained significantly more variance, 90.0% cumulatively, than expected by chance from an uncorrelated matrix with rank 7 (we used 9 traits, but *iWUE* and *LT* are linear combinations of other traits). We focus on the first principal component, denoted PC1, which explained a moderate amount of variation (35.5%). For reference, PC1 loaded positively with leaf thickness and *LMA*, but negatively with A_N_ and components of CO_2_ diffusion (*g*_m_, *g*_s_).

To test the hypothesis that species from dry and/or hot environments tradeoff efficient CO_2_ diffusion for a more robust, stress-tolerant leaf anatomy, we looked at the correlation between mean annual precipitation and temperature to species’ average position along PC1. For this analysis, we removed the three cultivars of *S. lycopersicum* var. *esculentum* and one accession *S. lycoperiscum* var. *cerasiforme*, an unimproved landrace. To reduce the influence of a single outlier species (*S. juglandifolium*), we shrank the interspecific variance of PC1 using a signed logarithm transformation:

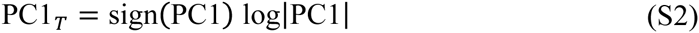

We also performed analyses with and without this species. We obtained mean annual precipitation and temperature at the habitat of origin for each source population from Worldclim (Hijmans *et al*. 2005), which has been used previously to study climatic adaptation in wild tomatoes (Chitwood *et al*. 2012; Nakazato *et al*. 2008). Phylogenetic regression was carried out with the R package phylolm version 2.2 (Ho & Ané 2014) using the ‘OUrandomRoot’ model to account for phylogenetic nonindependence. Other trait-climate relationships were tested in the same way.

## Results

### CO_2_ diffusion and biochemistry limit photosynthesis and alter water-use efficiency

Tomato species vary considerably in photosynthetic rate (*A*_N_) and intrinsic water-use efficiency (*iWUE*), driven by constraints on leaf CO_2_ diffusion and the maximum rate of carboxylation (*V*_cmax_). Phylogenetic linear mixed effects models (‘phyloLME’) showed that between individual plants, stomatal (*g*_s_) and mesophyll (*g*_m_) conductance increased *A*_N_ (Table 1; Figure 2A,B), but had opposing effects on *iWUE* (Table 1). However, phylogenetic relationship explained little of the trait variation, thus phylogenetic and nonphygenetic gave nearly identical results (Table 1). Increased g_s_ was associated with lower *iWUE* (Figure 2C), whereas greater *g*_m_ was associated with greater *iWUE* (Figure 2D). Like *g*_m_, greater biochemical capacity, as indicated by *V*_cmax_, was associated with significantly greater *A*_N_ and *iWUE* (Figure 2). Thus, species achieved high photosynthetic rates via two routes: high g_s_, but lower *iWUE* or high g_m_, but relatively high *iWUE*. The drawdown of CO_2_ concentration from the leaf interior (*C*_i_) to the chloroplast (*C*_c_), another indicator of diffusional constraint in the leaf mesophyll, was even more strongly correlated with *iWUE* than *g*_m_ (Figure S1), arguing that reduced internal diffusion enhances *iWUE*.

**Table 1:**
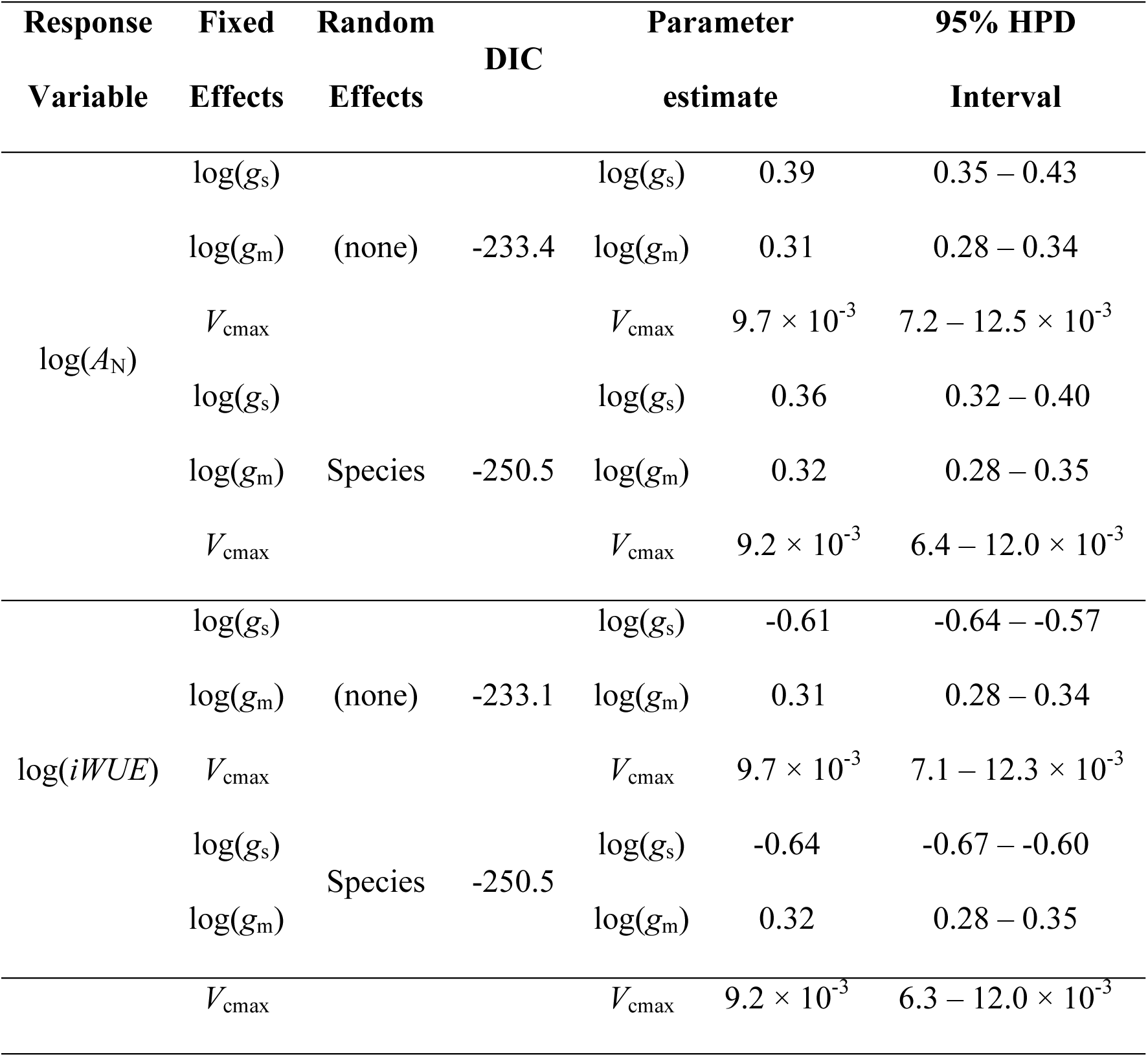
Diffusional and biochemical constraints drive variation in photosynthetic rate (*A*_N_) and intrinsic water-use efficiency (*iWUE*) among tomato species. For both *A*_N_ and *iWUE*, phylogenetic linear mixed effects models including ‘Species’ as a random effect improved model fit (lower Deviance Information Criterion [DIC]) over nonphylogenetic linear models, indicating some effect of phylogenetic relatedness. Greater stomatal (*g*_s_) and mesophyll (*g*_m_) conductance significantly increased *A*_N_ but had opposing effects on *iWUE*. Likewise, greater maximum carboxylation rates (*V*_cmax_) increased both *A*_N_ and *iWUE*. Parameters were estimated from the mode of 10^4^ samples drawn from the posterior distribution using MCMCglmm. The effects of *g*_s_, *g*_m_, and *V*_cmax_ are highly significant (95% high posterior density [HPD] intervals do not overlap zero) in all models.

**Figure 2:**
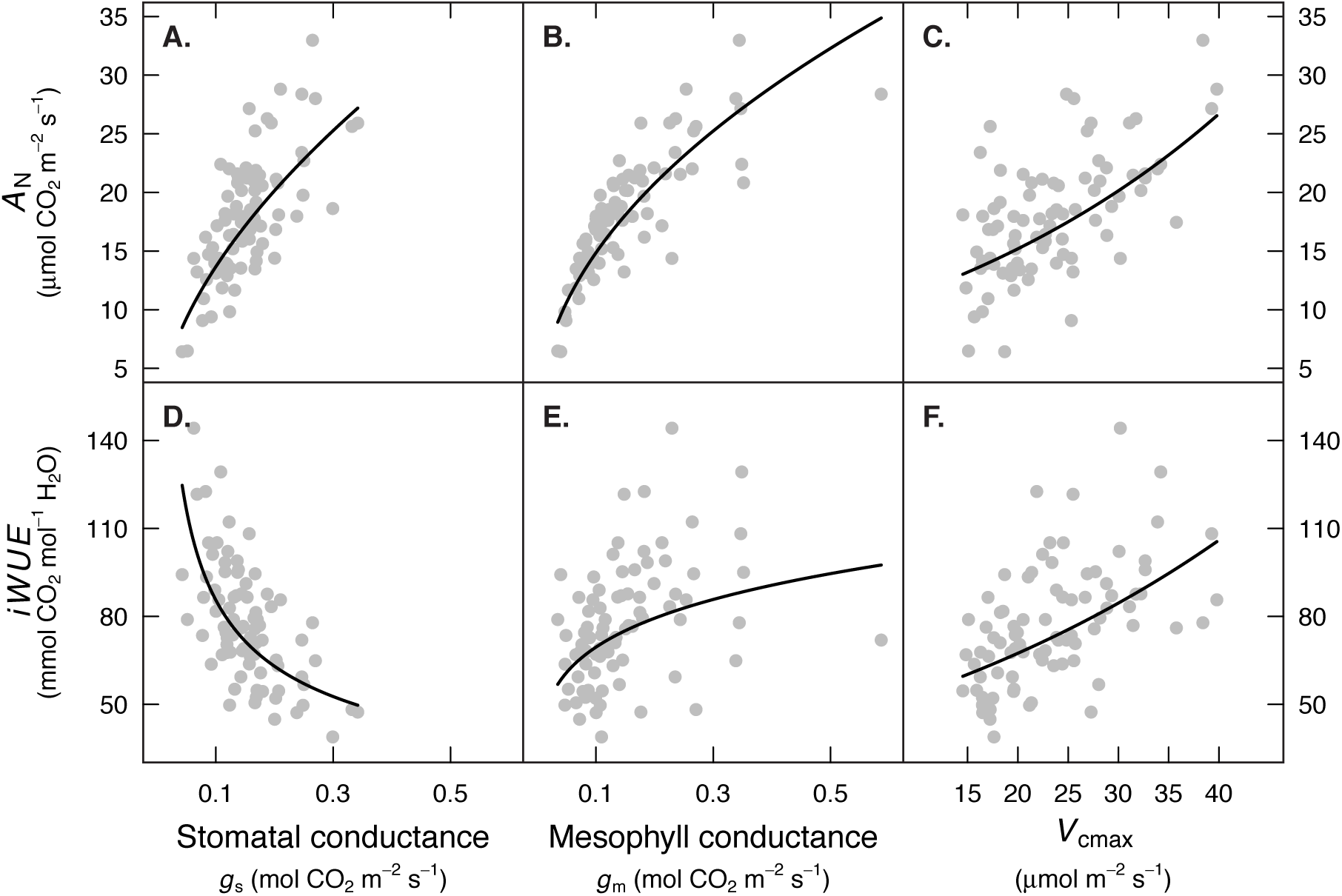
Leaf CO_2_ diffusion and biochemistry limit net CO_2_ assimilation rates (*A*_N_) and alter intrinsic water-use efficiency (*iWUE*). Each point is data from one of 82 individual plants across 19 wild and cultivated tomato taxa. Panels A. and B. show that faster diffusion through stomata (*g*_s_) and mesophyll (*g*_m_) resulted in higher *A*_N_. However, these parameters had opposing effects on *iWUE*. Greater *g*_s_ lowered *iWUE* (panel D.), whereas greater g_m_ increased *iWUE* (panel E.). Leaf biochemistry, specifically the maximum rate of carboxylation (*V*_cmax_) also increased *A*_N_ (panel C.) and *iWUE* (panel F.). Fitted lines are based on a Bayesian phylogenetic mixed models treating Species as a random effect: log(*A*_N_) ~ log(*g*_s_) + log(*g*_m_) + *V*_cmax_ + (1|Species) and log(*iWUE*) ~ log(*g*_s_) + log(*g*_m_) + *V*_cmax_ + (1|Species). For visual aid, lines were drawn to account for positive covariation between *g*_s_, *g*_m_, and *V*_cmax_ (data not shown). For example, panel A. shows the predicted *A*_N_ at a given *g*_s_ with *g*_m_ and *V*_cmax_ set to the predicted value at a given *g*_s_. All slopes were significantly different than zero (*P* < 0.0001) based on 10^4^ MCMC samples from the posterior distribution of the model.

### Limited variation in Rubisco biochemistry between tomato species

The two Rubisco LSu types showed significant differences in some of the kinetic parameters (Table 2; see Methods S1 for description of how types were identified and Results S1 for further detail on molecular evolution of *rbc*L in tomatoes). Interestingly, the Rubisco LSu type 2, which occurred in the domesticated clade species (*S. pimpinellifolium, S. cheesmaniae, S. galapagense, S. lycopersicum* var. *cerasiforme*, and the cultivars; see Figure S2) and *S. habrochaites*, had a higher *S*_c/o_ value, which was due to higher Michaelis-Menten constant for CO_2_ under atmospheric conditions (K_c_^air^) and lower catalytic turnover rate for the oxygenase reaction (k_cat_^o^) compared to the Rubisco LSu type 1. Non-significant differences were observed between the two Rubisco LSu types in the remaining kinetic parameters.

**Table 2.** Comparison of the Rubisco kinetics at 25°C for the two Rubisco LSu types detected among wild and domesticated tomatoes based on the amino acid sequence, namely LSu types 1 and 2. Parameters, measured in a representative species of each Rubisco type, describe the Michaelis-Menten constant for CO_2_ under 0% O_2_ (*K*_c_) and 21% O_2_ (*K*_c_^air^), the maximum rates of carboxylation (*k*_cat_^c^) and oxygenation (*k*_cat_^o^), carboxylation catalytic efficiency (*k*_cat_^c^/*K*_c_), and specificity factor (*S*_c/o_). Data are means ± SE of four replicates per Rubisco type. Significant differences between both Rubisco types are indicated by the ANOVA *P*-value, * <0.05; ** <0.01.

| Rubisco LSu type | 1 | 2 | $P$ value |
| --- | --- | --- | --- |
| Species measured | *S. lycopersicoides* | *S. lycopersicum* |  |
| $K_c$ (mM) | $9.3 \pm 0.9$ | $8.2 \pm 0.7$ | 0.308 |
| $K_c^{\text{air}}$ (mM) | $16.6 \pm 0.8$ | $13.5 \pm 0.8$ | 0.020* |
| $k_{\text{cat}}^c$ (s<sup>-1</sup>) | $2.430 \pm 0.138$ | $3.142 \pm 0.518$ | 0.176 |
| $k_{\text{cat}}^o$ (s<sup>-1</sup>) | $0.892 \pm 0.099$ | $1.359 \pm 0.108$ | 0.010** |
| $k_{\text{cat}}^c/K_c$ (s<sup>-1</sup> mM<sup>-1</sup>) | $0.261 \pm 0.026$ | $0.391 \pm 0.075$ | 0.127 |
| $S_{c/o}$ (mol mol<sup>-1</sup>) | $99.4 \pm 3.5$ | $115.0 \pm 2.1$ | 0.007** |

### Modest coordination between leaf physiological and structural traits

The first principal component (PC1) accounted for 35.5% of the variation among individual leaf physiology (*g*_m_, *g*_s_, *A*_N_, *V*_cmax_, *R*_dark_, *iWUE*) and bulk anatomy (*LMA, LT, LDMC*). On one end of this axis were thin leaves with fast CO_2_ diffusion and high *A*_N_; on the other were thick leaves with slower CO_2_ diffusion and lower *A*_N_ (Figure 3). This principal component indicates an axis of leaf trait variation likely mediated by tradeoffs between more robust leaf structure (i.e. higher *LMA* and *LT*) and CO_2_ diffusion. However, the modest amount of trait variance explained indicates that this tradeoff does not tightly constrain leaf trait evolution in tomatoes. The second principal component (PC2; 23.8% variance explained) was most strongly associated with *iWUE* and showed that greater *iWUE* was associated with higher water content (lower *LDMC*) leaves (Figure 3).

**Figure 3:**
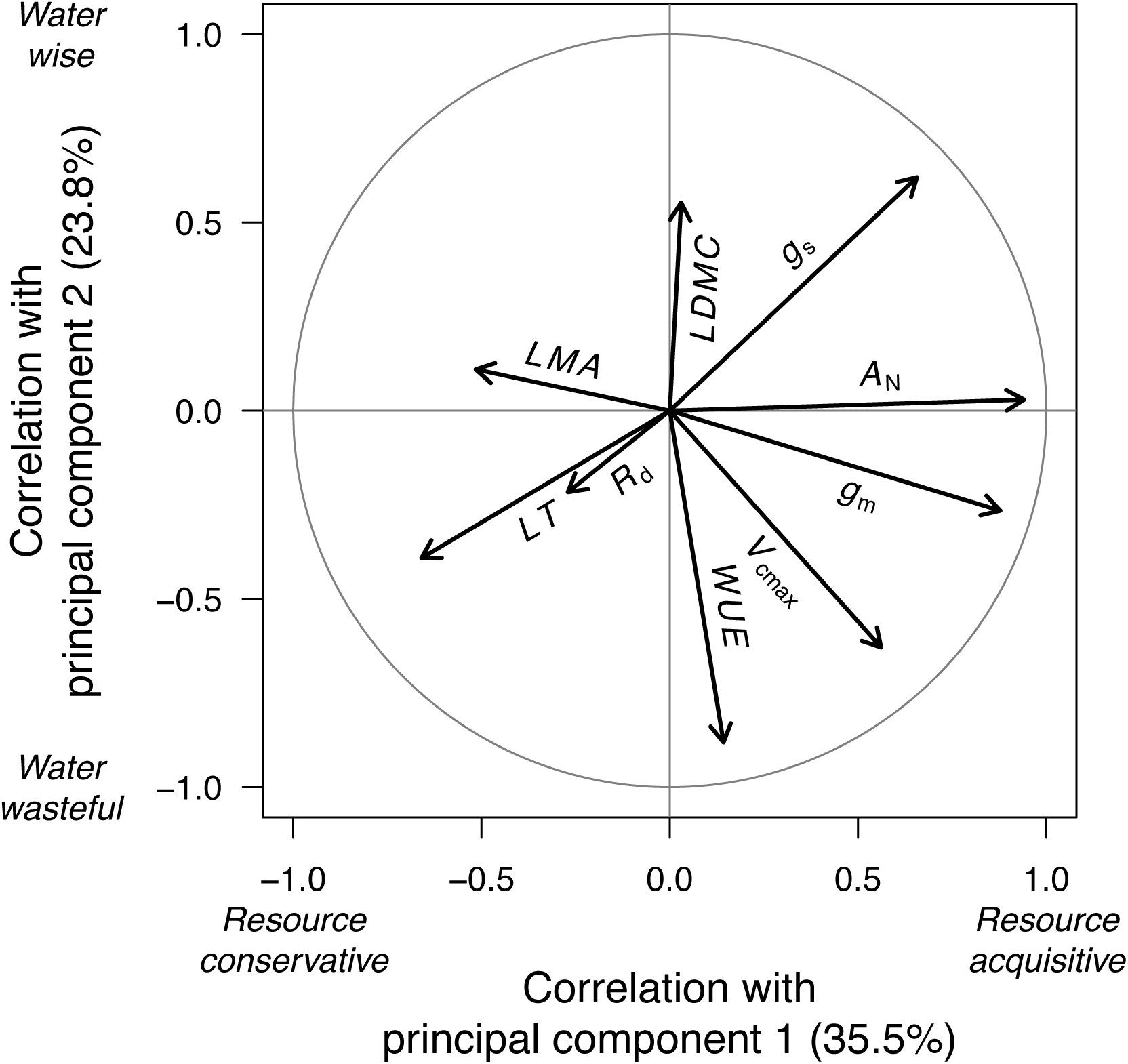
Major axes of leaf trait variation in tomato. The first axis, principal component 1, explains 35.5% of the total variation. It delineates plants along a continuum from resource acquisitive (higher PC1, greater *A*_N_, thinner leaves) to resource conservative (lower PC1, lower *A*_N_, thicker leaves). The second axis, principal component 2, explains 23.8% of the trait variation and delineates an axis from more water wise (higher PC2, greater *iWUE*) to water wasteful (lower PC2, lower *iWUE*). The arrows are vectors showing the correlation across individual plants between a trait and both principal components. For example, *A*_N_ is positively correlated with PC1, but uncorrelated with PC2. The grey circle shows the outer possible set of correlation combinations. Abbreviations: *R*_dark_ = dark respiration rate; *iWUE* = intrinsic water-use efficiency; *V*_cmax_ = maximum rate of carboxylation; *g*_m_ = mesophyll conductance; *A*_N_ = net CO_2_ assimilation rate; *g*_s_ = stomatal conductance; *LDMC* = leaf dry matter content; *LMA* = leaf mass per area; *LT* = leaf thickness.

### Limited evidence leaf trait-climate associations

Contrary to the hypothesis that dry, hot environments select for a stress-tolerant, robust leaf structure, thinner leaves and more rapid CO_2_ diffusion (i.e. higher values of PC1) was associated with drier habitats (PC1-Precip, *P* = 0.012), but not temperature (PC1-Temp, *P* = 0.069). However, this correlation between PC1 and precipitation was strongly influenced by a single species, *S. juglandifolium* (Figure 4), and was not significant if this species was removed (PC1-Precip, *P* = 0.266). Certain traits that loaded strongly with PC1, especially *LMA* and *LT*, were strongly associated with precipitation. Specifically, species from the driest habitats had the thinnest leaves, even when *S. juglandifolium* was excluded (Figure S3). Our data therefore do not support the hypothesis that species tradeoff slow CO_2_ diffusion and lower metabolic rates for robust leaf structure in stressful environments. All trends in the data are actually in the exact opposite direction.

**Figure 4:**
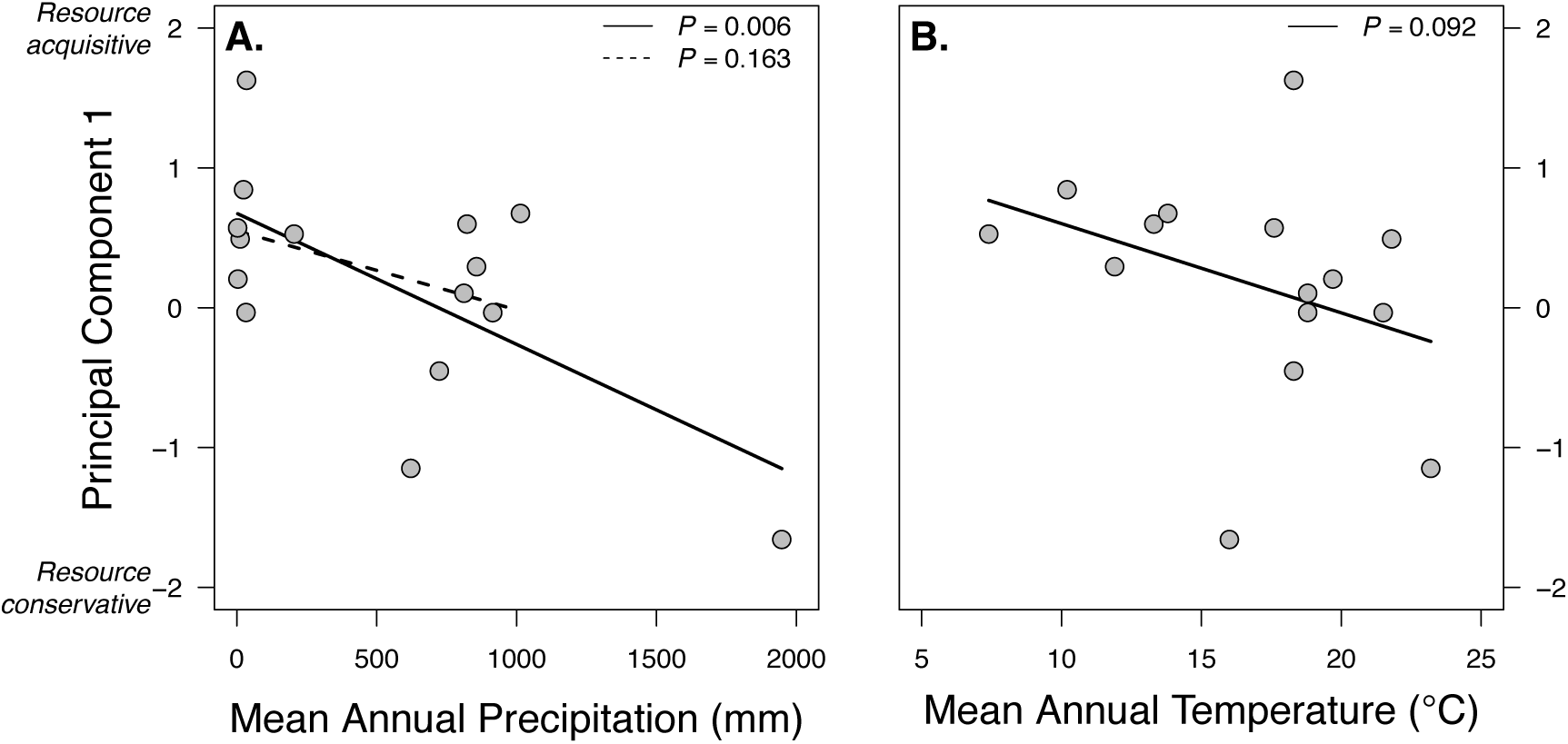
The resource conservative to acquisitive axis of leaf trait variation was uncorrelated with mean annual precipitation (A.) and temperature (B.). Each point shows the climate at habitat of origin for a different wild tomato taxon and its position, averaged from multiple individuals in the experiment, along the resource acquisitive to conservative axis. There was a significant positive correlation between PC1 and precipitation, but this was driven by a single influential species, *S. juglandifolium;* the correlation was not significant once this species was removed. All *P*-values are from phylogenetic linear regression.

## Discussion

Many ecologists seek to distill organismal variation down to a manageable number of key functional traits (e.g. Perez-Harguindeguy *et al*. 2013). Leaf traits in particular, which are necessarily constrained by the need to capture sunlight, CO_2_, and manage nutrient/water, may fall along a single ‘fast-slow’ continuum (Reich 2014 and references therein) associated with resource acquisitive and resource conservative traits (Mason & Donovan 2015). The key assumptions are that leaf structural traits such as leaf mass per area (*LMA*) strongly constrain photosynthesis by reducing CO_2_ diffusion and that there is strong selection for tight coordination between CO_2_ diffusion and biochemical capacity for photosynthesis. These assumptions have been evaluated at broad phylogenetic scales, but rarely addressed among closely-related species. This is important because patterns among distantly related species can be driven primarily by disparate functional groups (e.g. deciduous versus evergreen species) that have little bearing on the incremental evolutionary steps taken as species adapt to new environments.

In tomatoes, we found evidence for a ‘fast-slow’ spectrum mediated by diffusional/biochemical constraints and leaf structure, but this spectrum explained a modest amount of total leaf trait variation. There is no precise prediction for exactly how much coordination we *should* have seen under the coordination hypothesis, but variation was well-distributed across the first four components, indicating multiple important axes. Furthermore, although thicker leaves with greater *LMA* were associated with reduced CO_2_ diffusion, the relationships were weak (Figure S4), demonstrating ample scope for thick and/or dense leaves to have relatively high diffusion and *vice versa*. Finally, lack of coordination cannot be explained by relatively little physiological variation between species. Indeed, despite sharing a recent common ancestor, we observed a dramatic range in traits like *A*_N_ (6.4 − 33.0 μmol CO_2_ m^−2^ s^−1^) in wild tomatoes. In fact, our study should have been especially able to detect coordination, if it existed, because we measured plants with similar growth form and functional type in a common garden at the same age, eliminating sources of variation common in other studies. The lack of tight coordination between leaf structure, CO_2_ diffusion, and photosynthetic biochemistry means there may be multiple, loosely coordinated axes of leaf trait variation that provide a substrate for labile, unconstrained evolution in response to novel selective pressures from climate change and crop breeders.

In fact, many combinations of stomatal conductance (*g*_s_), mesophyll conductance (*g*_m_), and *V*_cmax_ resulted in similar photosynthetic rates but very different water-use efficiencies (Figure 2). Typically, we expect that to increase photosynthetic rate, plants must increase stomatal conductance, increasing transpirational loss and decreasing water-use efficiency. If mesophyll conductance and biochemical capacity were closely coordinated with stomatal conductance, then there would be limited opportunity to increase water-use efficiency without sacrificing photosynthetic rate. However, we find that there may be substantial scope to increase photosynthetic rate while maintaining high water-use efficiency. Indeed, photosynthetic rate and water-use efficiency were essentially uncorrelated. This axis of variation was evident in a second principal component that loaded positively with *iWUE* but was orthogonal to *A*_N_ (Figure 3). Why don’t all species have high photosynthetic rate and water-use efficiency? In nature, there are probably other tradeoffs, especially nitrogen limitation, that prevent species from having greater mesophyll conductance and/or *V*_cmax_.

Photosynthetic variation in tomatoes is primarily mediated by anatomical differences in leaves rather than differences in Rubisco kinetics. In contrast to other clades of C_3_ plants that have undergone rapid diversification into different environments, evolutionary changes in Rubisco kinetics (e.g. faster rates of carboxylation or greater affinity for CO_2_ over O_2_) do not appear to play a major role in the evolution of tomatoes. This contrasts with other plant groups like *Limonium* (Galmés *et al*. 2014), which underwent adaptive evolution of Rubisco kinetics during their radiations into novel environments. Future work is needed to understand why some clades respond to selection through changes in protein biochemistry, whereas groups like tomato seem to primarily differ in anatomical traits.

We also find no evidence that drier or hotter environments favor robust leaf structure or ‘slow’, resource conservative strategies (low conductance, low *V*_cmax_) in wild tomatoes (Figure 4). Obviously, our study does not have the statistical power of broad comparative analyses (e.g. Wright *et al*. 2005), but the trends in the data were not even in the predicted direction. If anything, there was a tendency for species from the driest habitats to be on the ‘fast’ end of the leaf trait spectrum (see also Easlon & Richards 2009), but this was largely influenced by a single species, *S. juglandifolium*. Thus, tomato species in dry habitats may rely primarily on a form of drought tolerance, growing fast when water is available and dying back or going dormant during droughts. Alternatively, leaf traits may be decoupled from other traits (e.g. root:shoot ratios) that confer alternative drought avoidance or tolerance mechanisms.

The modest coordination we observe between leaf structure and physiological function indicates that bulk structural traits like leaf mass per area are probably insufficient to identify the most important axes of trait variation (Figure S4). Traits like leaf thickness and leaf mass per area are probably important in setting the upper bounds on maximum potential CO_2_ diffusion and photosynthetic rate (Flexas *et al*. 2008), but realized values depend on the precise anatomical and biochemical features of the leaf that are not well-captured by bulk structural traits (Tosens *et al*. 2012; Tomás *et al*. 2013). This suggests the possibility that there are many unique axes that vary leaf anatomy and photosynthesis without large effects on traits like *LMA*. Each of these axes may be a different tradeoff that could be evolutionarily important in some taxa, but not others.

From the modest coordination between leaf structure and function in wild tomatoes, we can make three major conclusions:

1. The leaf functional traits most commonly used comparative ecology (e.g. *LMA*) may not be good proxies for leaf function, especially among closely-related species. This is not because there was little variation among tomatoes. Tomato leaves varied widely in diffusional, biochemical, and structural traits, but there was remarkably little covariation between bulk structure and function.
2. Hence, broad-scale patterns of functional trait covariation may not be useful in predicting adaptive evolution in the recent past or in response to climate change. Nor should we expect that the relationship between traits and environments observed at broad phylogenetic scales (e.g. thick leaves in hot, dry environments) will hold at smaller scales. If anything, we found that low-*LMA*, resource acquisitive traits were associated with dry environments, the opposite of what is commonly predicted (see also Mason & Donovan 2015).
3. Although our study challenges commonly-held assumptions about the relationships between leaf structure and function, this does not refute that intimate structure-function relationships exist. Rather, detailed anatomical traits such as guard cell dimensions, mesophyll structure, and photochemical enzyme concentrations are needed to identify the most important tradeoffs defining leaf anatomy and physiology.

Adaptive radiation of tomatoes and other plant groups undoubtedly requires evolution of new morphological and physiological traits, but this study suggests that we do not yet have a general explanation for variation in the most important leaf traits affecting photosynthesis.

## Acknowledgements

CDM was supported by an Evo-Devo-Eco Network (EDEN) research exchange (NSF IOS #0955517). The research was supported by project AGL2013–42364-R (Plan Nacional, Spain) awarded to JG. We acknowledge Trinidad García at the radioisotope service (UIB), and Miquel Truyols and collaborators of the UIB Experimental Field and Greenhouses for their technical support. Chase Mason provided feedback.

---

**Figure S1:**
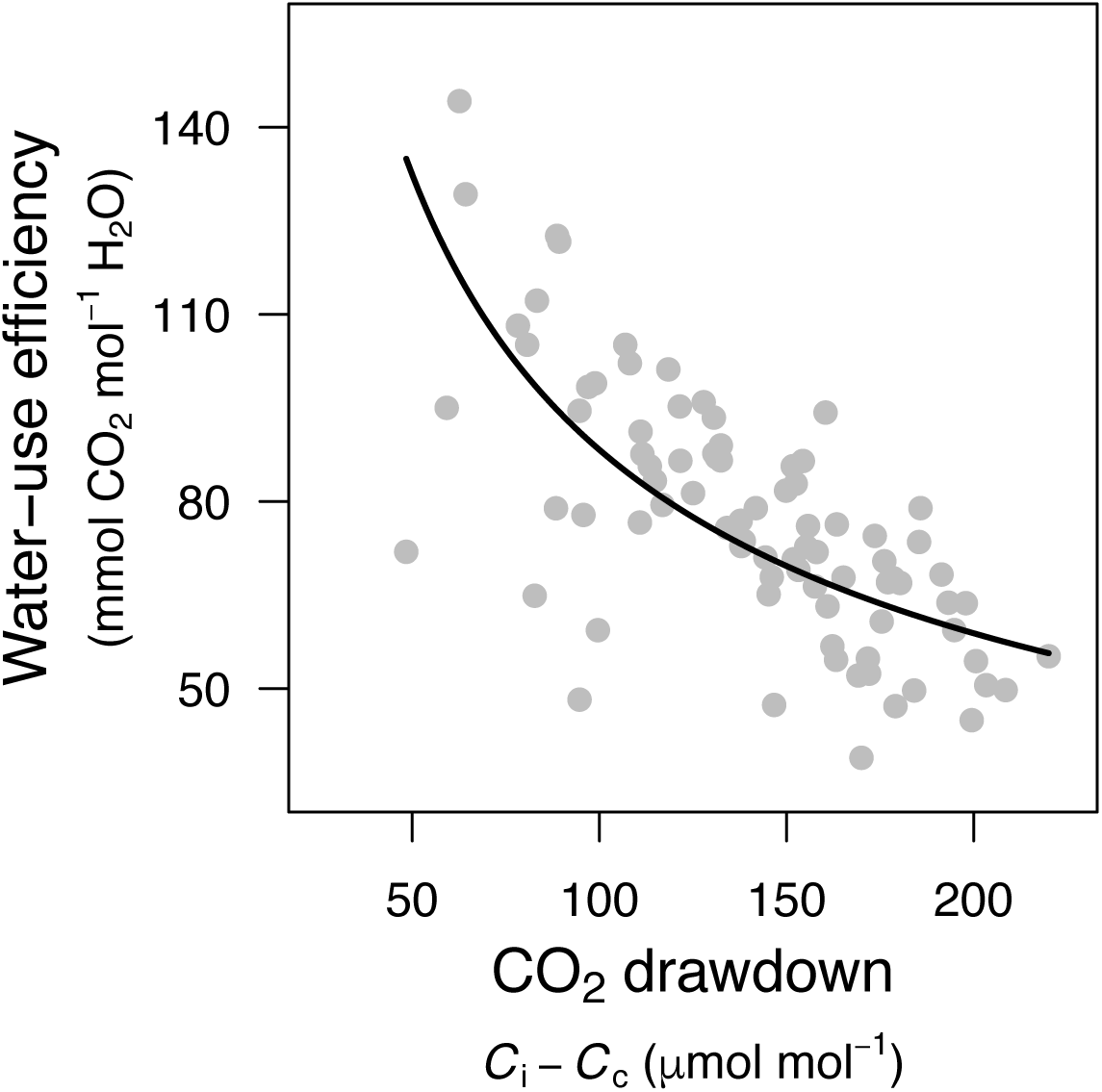
CO_2_ drawdown from the leaf interior (*C*_i_) to the chloroplast (*C*_c_) is negatively associated with i*WUE* in tomatoes, indicating that reduced diffusional constraints (lower *C*_i_ – *C*_c_) leads to increased *iWUE*. Fitted line is based on a Bayesian phylogenetic mixed models treating Species as a random effect: log(*iWUE*) ~ log(*C*_i_ − *C*_c_) + (1|Species). The slope is significantly different than zero (*P* < 0.0001) based on 10^4^ MCMC samples from the posterior distribution of the model.

**Figure S2:**
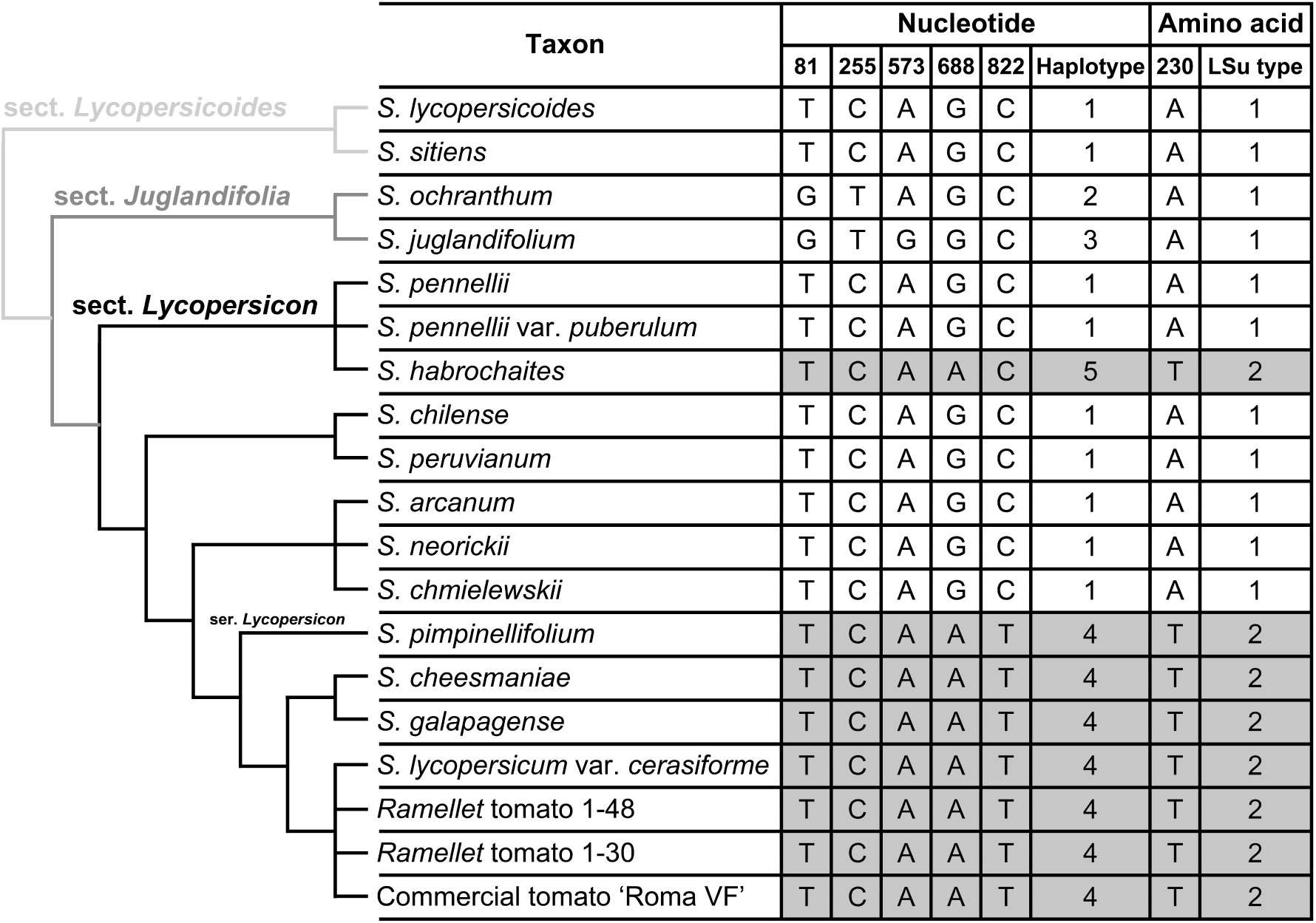
Evolutionary patterns of the Rubisco large subunit (LSu) among wild and domesticated tomato species. The species relationships shown are based on an 18-gene phylogeny from Haak *et al*. (2014; see Methods). Nucleotide differences in the *rbc*L gene are indicated, giving rise to five different haplotypes. Those mutations result in a single difference at amino acid level and thus two Rubisco LSu types within tomatoes.

**Figure S3:**
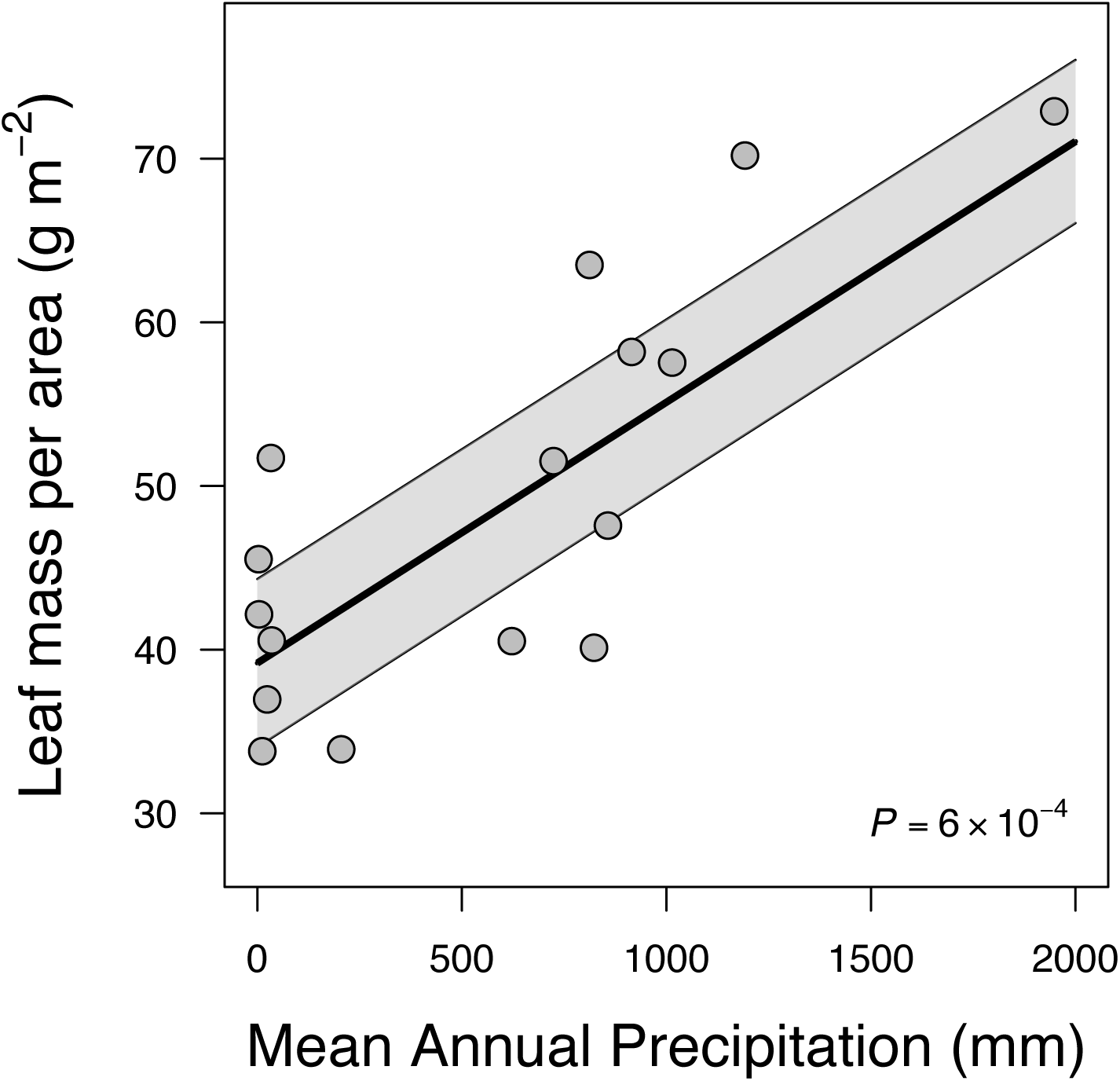
Leaf mass per area (*LMA*) was positively correlated with mean annual precipitation. Each point shows the climate at habitat of origin for a different wild tomato taxon and its *LMA*, averaged from multiple individuals in the experiment. There was a significant positive correlation between PC1 and precipitation. All *P*-values are from phylogenetic linear regression.

**Figure S4:**
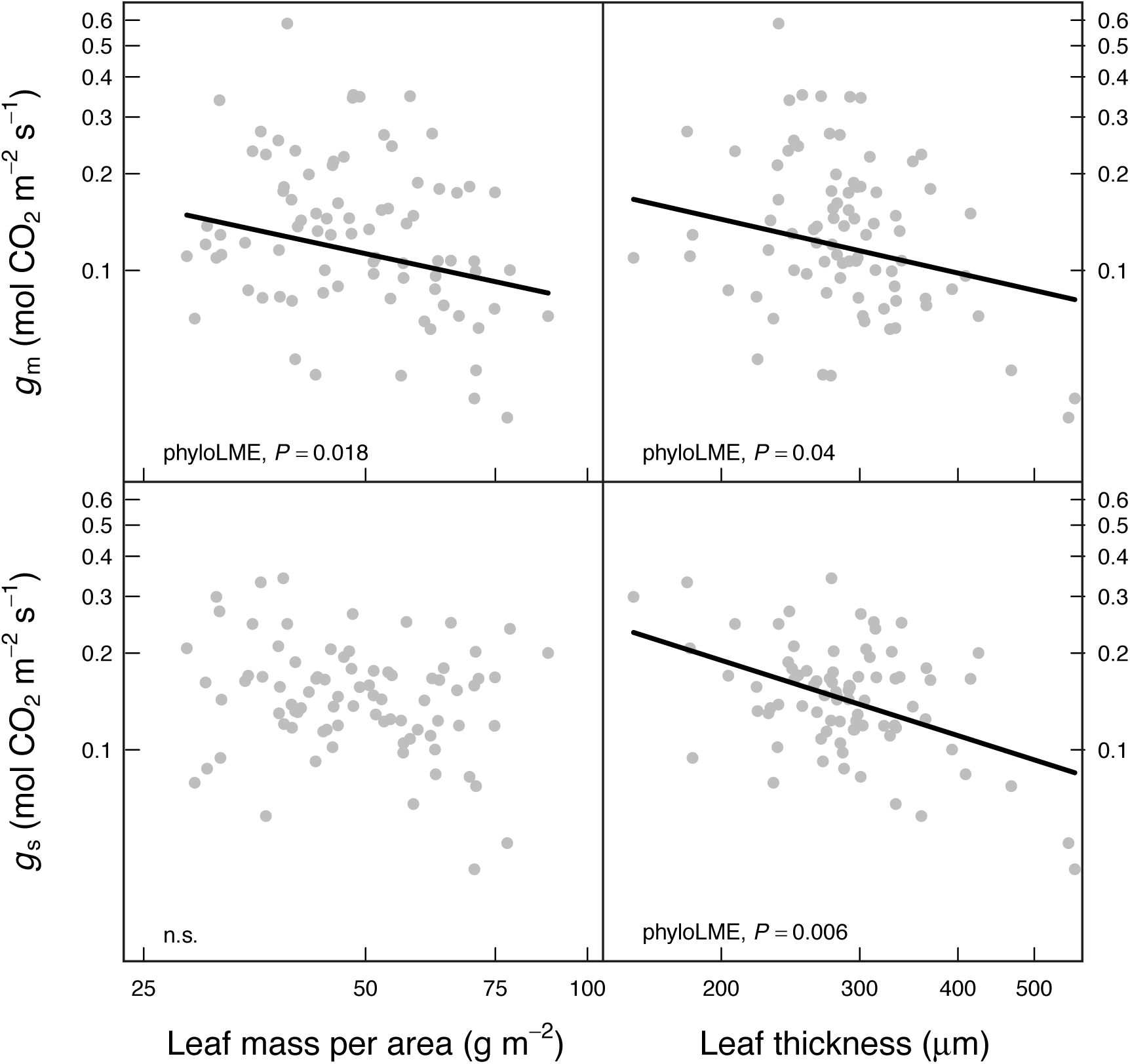
Bulk leaf structure (leaf mass per area [*LMA*] and leaf thickness [*LT*]) weakly constrain leaf CO_2_ diffusive conductance through stomata (*g*_s_) and mesophyll (*g*_s_). Each point is data from one of 80 individual plants across 19 wild and cultivated tomato taxa. Panels A. and B. show that diffusion through mesophyll (*g*_m_, mesophyll conductance) was limited by higher *LMA* and *LT*. Additionally, *LT* (panel D.), but not *LMA* (Panel C.) were associated with decreased stomatal conductance (*g*_s_) parameters. Fitted lines are based on Bayesian phylogenetic mixed models treating Species as a random effect. Mixed models with *LT* also included *LDMC*, but this did not have a significant effect (results not shown). All *P*-values are based on 10^4^ MCMC samples from the posterior distribution of the model.

## Methods S1

### Gas exchange

We used an open path infrared gas exchange analyzer with a 2-cm^2^ leaf chamber fluorometer (LI-6400–40, LI-COR Inc., Lincoln, NE, USA) to simultaneously measure leaf gas exchange and chlorophyll *a* fluorescence. To minimize leaf position and age effects, all measurements were made on young, fully-expanded leaves (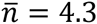, range = [3, 5]). The ambient CO_2_ concentration in the chamber (*C*_a_) was 400 μmol CO_2_ mol^−1^ air, leaf temperature was 25° C, photosynthetic photon flux density (PPFD) was 1500 μmol m^−2^ s^−1^ with 90:10 red:blue light, and relative humidity was between 40 − 60%. Once a leaf reached steady-state photosynthesis (*A*) and stomatal conductance (*g*_s_), usually after ~30 min, we measured the response to changing substomatal CO_2_ concentrations (*C*_i_) by adjusting the ambient CO_2_ concentration in the leaf chamber (*C*_a_). We used 11 concentrations between 0 and 1750 μmol CO_2_ mol^−1^ air. The flow rate was 300 μmol s^−1^. Diffusional leaks for CO_2_ were corrected for using the methods of Rodeghiero *et al*. (2007). We observed no differences between diffusion coefficients calculated for leaves of different species or an empty chamber (data not shown).

From fluorescence measurements and *A* − C_i_ curves, we determined the net CO_2_ assimilation rate at *C*_a_ = 400 μmol CO_2_ mol^−1^ air (*A*_N_), stomatal conductance (*g_s_*), mesophyll conductance to CO_2_ (*g_m_*), and the maximum rate of carboxylation (*V_cmax_*). For all analyses, we used *g*_s_ and *g*_m_ at ambient *C*_a_ (400 +/− 5 μmol CO_2_ mol^−1^ air) because these values are most ecologically relevant and allowed us to directly analyze how *g_s_* and *g_m_* limited *A*_N_. Photosynthetic rate and stomatal conductance are estimated directly from gas exchange measurements. We estimated mesophyll conductance using from the equation (Harley *et al*. 1992):

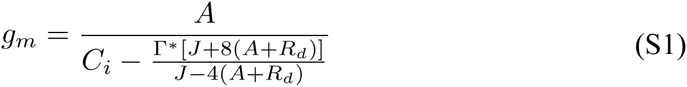

We measured the mitochondrial respiration rate (*R*_dark_) for each plant at predawn and used the common assumption *R*_d_ = *R*_dark_ / 2. As described below, we measured the chloroplastic CO_2_ compensation point (*Γ*^*^) *in vitro* from two species, *S. lycopersicoides* and *S. lycopersicum* var. *esculentum* cv. ‘Roma’, one each from the two LSu amino acid sequences among tomato species (see Results). For *S. lycopersicoides* (Rubisco LSu Type 1, Figure S2), we estimated *Γ^*^* = 40.46 μmol CO_2_ mol^−1^ air; for *S. lycopersicum* (Rubisco LSu Type 2, Figure S2) we estimated *Γ^*^* = 35.39 μmol CO_2_ mol^−1^ air. These values were applied to the other species with the same Rubisco LSu Type (Figure S2).

The electron transport rate of photosystem II (*J*_f_) was calculated from fluorescence measurements as *J*_f_ = Φ_PSII_*Iαβ*, where Φ_ΡSII_ is the quantum yield (moles of CO_2_ fixed per mole of quanta absorbed) of photosystem II, *I* is irradiance, α is the leaf light absorptance, and β is the photosystem partitioning factor. Φ_ΡSII_ was estimated from the chlorophyll fluorescence data. We estimated the product αβ separately for each species using the relationship 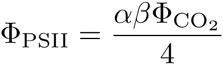 under nonphotorespiratory conditions (Genty, Briantis & Baker, 1989). To vary the quantum yield, we measured photosynthetic light response curves under 2% O_2_. In summary, we incorporated data on species-specific respiration (*R*_d_), Rubisco kinetics (*Γ*^*^), and light absorptance/photosystem partitioning (αβ) to improve the accuracy of *g_m_* estimates.

Using the calculated values of *g*_m_, we used the relationship:

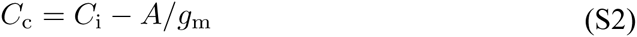

to determine *A* − *C*_c_ curves to estimate the maximum velocity of carboxylation (*V_c_*_max_ [μmol m^−2^ s^−1^]), which is proportional to the concentration of activated Rubisco. We estimated *V_c_*_max_ following Long & Bernacchi (2003) from the linear, Rubisco limited portion of the *A* − *C_c_* curve, empirically determined to be *C*_i_ < 200 mol CO_2_ mol^−1^ air. We used kinetic values from Table 2 (methods described below) to calculate effective Michaelis-Menten constants for each LSu type.

### Sequencing the Rubisco large subunit

Total DNA was extracted from fresh leaf tissue using the DNeasy Plant MiniKit (Qiagen). The *rbc*L gene coding for the Rubisco large subunit (LSu) was amplified using primers specifically designed for *Solanum*, the forward primer SoSpaF (5′-ATGAGTTCTAGGGAGGGAT-3′) and the reverse primer So1462R: (5’-GCAGGAAATAAAGAAGGATAAGG-3’). The BioMix Red reaction mix (Bioline Ltd., London, UK) was used to carry out the polymerase chain reaction (PCR) with the following conditions: an initial cycle of 95 °C, 2 min; 55 °C, 30 s; 72 °C, 4 min, followed by 36 cycles of 93 °C, 30 s; 53 °C, 30 s; 72 °C, 3.5 min. PCR products were visualized on 1% agarose gels, purified using the High Pure PCR Product Purification Kit (Roche, Germany) and sequenced with an ABI 3130 Genetic analyzer using the ABI BigDyeTM Terminator Cycle Sequencing Ready Reaction Kit (Applied Biosystems, Foster City, California). Due to the *rbc*L length, sequences were obtained in a two-fragment fashion, using the primer So1462R and the primer rbcL638R (5’-CGCATAAATGGTTGGGAATTC-3’) (Chen 1998). Sequence chromatograms were checked and corrected with the Chromas software and the contigs were assembled using BioEdit v7.1.3 software (Hall 1999). MEGA 5 (Tamura *et al*. 2011) was used to convert DNA to amino acid, and to align both sequence types.

### Rubisco catalytic characterization

Based on the two different amino acid sequences detected, a representative species for each type was selected, *S. lycopersicoides* for Rubisco LSu Type 1, and a domesticated accession *S. lycopersicum* var. *esculentum* cv. ‘Roma’ for the Rubisco LSu Type 2. We characterized Rubisco kinetics following Galmés *et al*. (2014). Rates of Rubisco ^14^CO_2_ fixation using fresh leaf protein extract were measured in 7 ml septum-capped scintillation vials, containing reaction buffer (yielding final concentrations of 100 mM Bicine-NaOH, pH 8.0, 20 mM MgCl_2_, 0.4 mM RuBP and ca. 100 W-A units of carbonic anhydrase) and one of nine different concentrations of CO_2_ (0–80 μM, each with a specific radioactivity of 3.7 × 10^10^ Bq mol^−1^), each at two concentrations of O_2_ (0 and 21% v/v), as described previously (Parry *et al*. 2007). Assays (1.0 ml total volume) were started by the addition of activated leaf extract, and the maximum velocity of carboxylase activity (*V*_max_) together with the Michaelis-Menten constant (*K*_m_) for CO_2_ (*K*_c_) determined from the fitted data. The K_m_ for the oxygenase activity was calculated from the relationship *K*_c,(21%O2)_ = *K*_c,(0%O2)_ · (1 + [O_2_]/*K*_o_). The [O_2_] was assumed to be 265 μM, but corrected for partial pressure by taking account of the atmospheric pressure and water saturated vapor pressure. Replicate measurements (*n* = 4) were made using protein preparations from different leaves of different individuals. For each sample, the maximum rate of carboxylation (*k*^c^_cat_) was extrapolated from the corresponding *V*_max_ value after allowance was made for the Rubisco active site concentration, as determined by [^14^C]CPBP binding (Yokota & Canvin 1985). Rubisco CO_2_/O_2_ specificity (*S*_c/o_) was measured as described previously (Galmés *et al*. 2005) using enzyme purified by polyethylene glycol (PEG) precipitation and ion exchange chromatography, and the values given for each species were the mean of 7–8 repeated determinations. The maximum oxygenation rate (*k*^o^_cat_) was calculated using the equation *S*_c/o_ = (*k*^c^_cat_ / *K*_c_)/(*k*^o^_cat_ / *K*_o_). All kinetic measurements were performed at 25°C.

## Results S1: Sequence comparisons reveal two Rubisco LSu types in tomatoes

We identified five mutations in the *rbc*L gene sequence (Figure S2), encoding for the Rubisco LSu. However, only one of these five mutations altered the amino acid sequence. The limited variability in LSu among tomato species contrasts with previous studies reporting large variability among closely related taxa in *Limonium* (Galmés *et al*. 2014), *Schidea* (Kapralov & Filatov 2006), *Amanthaceae* (Kapralov *et al*. 2012), and *Quercus* (Hermida-Carrera *et al*. unpub. data). The single mutation leading to a different amino acid sequence for the LSu at position 230 (Figure S2) resulted in improved *S*_c/o_ (i.e., higher specificity of the enzyme for CO_2_), but did not reduce the enzyme velocity (*k*^c^_cat_; Table 2). The same amino acid replacement A230 to T230 was observed in the Hawaiian endemic genus *Schiedea* (Kapralov & Filatov 2006). *Schiedea* represents one of the largest plant adaptive radiations on Hawaii and, similarly to the wild species of tomato in the present study, comprises closely related species with a broad range of morphological and ecological forms, inhabiting diverse environments from rainforest to desert-like conditions. According to Kellogg & Juliano (1997), residue 230 interacts with the *β*Α-*β*Β loop of small subunit, and replacement A230 to T230 causes a decrease in the hydrophobicity of the residue with effects in the stability at this position.

We designate the sequence associated with lower S_c/o_ ‘Rubisco LSu Type 1’ and that with higher *S*_c/o_ ‘Rubisco LSu Type 2’. Interestingly, the Rubisco LSu Type 2 occurs in the closest wild relatives of the domesticated tomato (i.e., from here on “the domesticated clade”: *S. pimpinellifolium, S. cheesmaniae, S. galapagense, S. lycopersicum* var. *cerasiforme*, and the domesticated accessions; see Figure S2), maybe pointing to this biochemical improvement at Rubisco level to one of the reasons triggering the diversification of the most recent tomato species, and from which the domesticated tomato was originated. Also, the lack of DNA mutations in the *rbc*L among those species could reflect a very recent occurrence of this Rubisco LSu type.

The phylogenetic relationship among sections *Lycopersicoides, Juglandifolia* and *Lycopersicon* is consistent both based on nuclear markers and cpDNA markers (Peralta *et al*. 2008; Robertson *et al*. 2011), and indicates that sect. *Lycopersicoides* diverged before sect. *Juglandifolia* (Figure 1). Therefore, since the cpDNA of sect. *Lycopersicon* is identical to that of sect. *Lycopersicoides* but different to that of sect. *Juglandifolia*, mutations in the latter section may have occurred after the split from sect. *Lycopersicon*, which maintained the original sect. *Lycopersicoides* cpDNA. Therefore, sect. *Juglandifolia* shows two exclusive mutations (in positions 81 and 255), and within *S. juglandifolium* has a third one (in position 573; Figure S2), which shows that this section has the most divergent *rbc*L sequences among tomatoes. However, these mutations did not alter the amino acid sequence of the LSu, suggesting that negative selection on the majority of nonsynonymous substitutions.

As an exception to the above is S. *habrochaites*, which diverged early on in the sect. *Lycopersicon* clade with *S. pennellii*. This species has the Rubisco LSu Type 2, but a different, exclusive haplotype as compared to that in the domesticated clade. Therefore, it appears that the mutation in the DNA position 688, resulting in the amino acid change leading to Rubisco LSu Type 2, may have occurred independently in *S. habrochaites* and in the domesticated clade species. Evidence for convergence is reinforced by the mutation in the DNA position 822, which occurs exclusively in all the domesticated clade species, whereas the *S. habrochaites* haplotype resembles other species in the Rubisco LSu Type 1 at this position. This apparently convergent origin of the mutation leading to an improved Rubisco would reflect 1) the importance of maintaining the ancestral (i.e., Type 1, sect. *Lycopersicoides*) Rubisco LSu across the tomato radiation, because it has been maintained in most species, and 2) little positive selection on the LSu, because only a single amino acid change was detected, in the amino acid position 230, and moreover, it may have occurred twice.

